# PERADIGM: Phenotype Embedding Similarity-based Rare Disease Gene Mapping

**DOI:** 10.1101/2025.04.01.646670

**Authors:** Wangjie Zheng, Yuhan Xie, Jianlei Gu, Hongyu Li, Stefan Somlo, Whitney Besse, Hongyu Zhao

## Abstract

Identifying genes associated with rare diseases remains challenging due to the scarcity of patients and the limited statistical power of traditional association methods. Here, we introduce PERADIGM (**P**henotype **E**mbedding similarity-based **RA**re **DI**sease **G**ene **M**apping), a novel framework that leverages natural language processing techniques to integrate comprehensive phenotype information from electronic health records for rare disease gene discovery. PERADIGM employs an embedding model to capture relationships between ICD-10 codes, providing a nuanced representation of individual phenotypes. By utilizing patient similarity scores, it enhances the identification of candidate genes associated with disease-specific phenotypes, surpassing conventional methods that rely on binary disease status. We applied PERADIGM to the UK Biobank dataset for three rare diseases: autosomal dominant polycystic kidney disease (ADPKD), Marfan syndrome, and neurofibromatosis type 1 (NF1). PERADIGM identified additional candidate genes associated with ADPKD-related and Marfan syndrome-related phenotypes, some of which are supported by existing literature, and demonstrated enhanced signal detection for NF1-specific phenotypes beyond traditional methods. Our findings demonstrate the potential of PERADIGM to identify genes associated with rare diseases and related phenotypes by incorporating phenotype embeddings and patient similarity, providing a powerful tool for precision medicine and a deeper understanding of rare disease genetics and clinical manifestations.

## 1 Introduction

The advent of Whole Exome Sequencing (WES) and Whole Genome Sequencing (WGS) [1] has facilitated the discovery of rare variants and their associations with both common and rare diseases. Numerous rare variant association methods have been developed to analyze WES and WGS data [2], characterizing a wide range of genotype-phenotype relationships. Despite some successes, these methods lack statistical power when the target phenotype is rare, which is often the case for rare diseases [3], even in large biobanks. Moreover, because rare diseases are often driven by rare variants, the likelihood of detecting significant associations is further reduced. For example, in the UK Biobank [4], which includes over 500,000 participants, the number of individuals with specific rare diseases is often small, making it difficult to identify associations using traditional rare variant association methods such as the Burden test [5], SKAT [6], and SKAT-O [7]. Historically, research on Mendelian diseases has focused on pathogenic variants. For instance, Autosomal Dominant Polycystic Kidney Disease (ADPKD) is primarily attributed to pathogenic variants in the *PKD1* and *PKD2* genes [8], while Marfan syndrome [9] and Neurofibromatosis type 1 (NF1) [10] are mainly associated with the *FBN1* and *NF1* genes, respectively. However, known pathogenic variants explain only a fraction of disease cases, suggesting the presence of additional candidate genes associated with disease-specific phenotypes. Identifying these genes remains challenging for traditional rare variant association methods due to the limited number of rare disease patients, even within large biobank cohorts.

Rare disease patients typically present unique phenotypic profiles [11], largely due to the strong effects of rare pathogenic variants [12]. As a result, these patients often exhibit distinctive phenotypes beyond their primary disease status. However, current methods primarily rely on binary disease classifications, overlooking the rich phenotype data available. Electronic Health Records (EHR) [13] provide a valuable source of comprehensive phenotype information across large cohorts. In the UK Biobank inpatient dataset [4], longitudinal phenotype data for each individual are recorded using the 10th revision of the International Classification of Diseases (ICD-10) codes [14]. Most studies incorporate ICD-10 phenotype information as binary variables in association analyses, including genome-wide association studies (GWAS) [15], phenotype-wide association studies (PheWAS) [16], and rare variant association tests [6, 7, 17]. However, treating ICD-10 codes as binary variables may lead to information loss, particularly in capturing relationships among phenotypes. Relevant secondary manifestations may be present even when the primary diagnosis is not explicitly coded, either due to incomplete clinical documentation or enrollment prior to a formal diagnosis. While this may be less common in the UK Biobank cohort, which primarily includes older individuals, it remains a possibility. Additionally, for some rare diseases, precise ICD-10 codes may be missing in the UK Biobank because participants tend to represent a relatively healthy population, and such diagnoses may be underreported or undetected. Leveraging the full spectrum of phenotype information can therefore provide a more comprehensive view of disease expression. In ADPKD, while *PKD1* and *PKD2* mutations are known to cause both kidney and liver cysts, the presence of liver cysts without kidney involvement may indicate other genetic causes. More broadly, the reliance on discrete ICD-10 codes can obscure phenotypic nuances, potentially limiting the identification of additional candidate genes associated with rare disease phenotypes. Besides identifying candidate genes for rare diseases, it is also valuable to uncover genes associated with disease-related phenotypes using EHR data. These phenotype-associated genes may provide further insights into genetic modifiers of disease phenotypes, secondary complications, and potential therapeutic targets. Addressing the limitations of ICD-10-based approaches by incorporating richer phenotype representations could improve the power of rare disease gene discovery and enhance our understanding of genotype-phenotype relationships.

To enhance phenotype information for candidate gene discovery, researchers increasingly apply natural language processing (NLP) models to phenotype data [18]. These models transform EHR phenotype terms into high-dimensional vector representations, generating phenotype embeddings that synthesize the entirety of available clinical data. Unlike traditional approaches that rely solely on ICD-10 codes, NLP-based embeddings can capture richer phenotypic representations by incorporating contextual information from structured and unstructured medical records. Established NLP embedding models, such as Word2Vec [19], generate static embeddings for medical events based on their co-occurrence patterns and relationships within the data. This approach enables phenotype embeddings to encode both the presence of specific phenotypes and their relationships with other clinical features documented in EHR, providing a more nuanced representation of a patient’s condition. By integrating textual and coded clinical data, NLP-driven embeddings offer a promising alternative to binary disease status representations, which may overlook critical phenotypic complexity. While some studies [20, 21] have incorporated embedding models in gene discovery, their applications to rare diseases remain limited due to the scarcity of disease samples. As a result, few methods have leveraged embedding models for identifying candidate genes associated with rare disease phenotypes. Expanding NLP-based approaches in rare disease research could provide a more comprehensive understanding of genotype-phenotype relationships and improve the identification of relevant genetic contributors.

To address this research gap, we have developed a principled framework called **P**henotype **E**mbedding similarity-based **RA**re **DI**sease **G**ene **M**apping (**PERADIGM**) that integrates genetic data with phenotype information derived from ICD-10 codes. Using an embedding model to represent each individual’s phenotype profile, we found that rare disease patients and loss-of-function variant carriers exhibit significantly higher phenotypic similarity within their respective groups compared to random controls. Leveraging this similarity, we expanded the effective pool of rare disease patients, thereby increasing statistical power for identifying candidate genes associated with disease-specific phenotypes. We evaluated PERADIGM on three rare diseases: ADPKD, Marfan syndrome, and *NF1* using data from the UK Biobank. By systematically scanning all genes, PERADIGM identified significant candidate disease-modifying genes in which rare variants may affect the disease-related phenotypic spectrum, uncovering additional associations beyond those detected by traditional methods, some of which are supported by prior studies. These findings demonstrate that PERADIGM can effectively extract and utilize phenotype information through embedding models, leveraging patient similarity to improve the identification of candidate disease-modifying genes for rare diseases.

## 2 Results

### 2.1 Overview of PERADIGM

PERADIGM is a phenotype embedding similarity-based statistical framework designed to identify candidate genes associated with disease-specific phenotypes for rare diseases using ICD-10 codes and sequencing data. It aims to improve statistical power in gene discovery, particularly in biobank datasets where some cases may be missed or misclassified during diagnosis. In this context, we define “risk genes” as candidate genes whose rare variants are associated with phenotypic manifestations related to the target disease. Importantly, investigating genes associated with disease-specific phenotypes, rather than focusing solely on the disease diagnosis itself, may provide additional insights into the underlying mechanisms of disease progression, variability in clinical presentation, and potential genotype-phenotype relationships. The PERADIGM framework employs a two-stage procedure, as detailed below.

In the first stage, we utilize individual-level ICD-10 codes from EHR data to create embedding representations for each phenotype using a continuous bag-of-words (CBOW) Word2Vec model. Patient embeddings are then generated by taking a weighted average of the phenotype embeddings, where the weights are determined based on two considerations. First, we consider the statistical evidence of association between each phenotype and the target disease. Second, we account for the prevalence of each phenotype in the biobank. Further details are provided in the Methods section. This weighting strategy captures both the phenotype’s relevance to the target disease and its information content, resulting in an embedded phenotype profile for each individual.

In the second stage, we conduct a genome-wide scan to calculate a risk score for each candidate gene with respect to the target rare disease, based on the embedded individual profiles. We hypothesize that genes associated with the target disease-associated phenotypes will show greater phenotypic similarity between individuals carrying rare loss-of-function (LoF) variants and those diagnosed with the disease. In PERADIGM, the similarity between two individuals is measured using the cosine similarity of their embedded phenotype profile vectors. We then derive a gene-specific risk score by comparing the phenotype embedding similarity between individuals carrying rare LoF variants and individuals diagnosed with the target disease. A higher similarity—and thus a higher risk score—between LoF variant carriers and disease cases suggests a higher likelihood that the candidate gene is associated with the disease. We assess the statistical significance of the observed score by comparing it to an empirical null distribution generated from repeated random sampling of individuals without regard to carrier status.

We note that PERADIGM uses widely available ICD-10 codes to identify candidate genes associated with rare disease-asscociated phenotypes, while some existing methods [22–24] integrate phenotype similarity using Human Phenotype Ontology (HPO) terms. By leveraging ICD-10 codes, PERADIGM can be more easily applied across diverse datasets and large-scale biobanks. In the following sections, we demonstrate the utility of PERADIGM through its application to several rare diseases using the first 200K WES release from the UK Biobank.

### 2.2 ADPKD

We first applied PERADIGM to identify candidate genes associated with ADPKD-related phenotypes [25]. ADPKD is one of the most common inherited kidney disorders, affecting approximately 1 in 400 to 1 in 1,000 individuals. It is characterized by the development of numerous fluid-filled cysts in the kidneys, leading to progressive renal dysfunction and, eventually, kidney failure. Approximately 78% of ADPKD cases are caused by pathogenic variants in the *PKD1* gene, while another 15% result from variants in the *PKD2* gene [26]. These genes encode polycystin-1 and polycystin-2, respectively, which are essential for maintaining the structural integrity of renal tubular cells and regulating calcium signaling. Disruptions to these proteins impair normal cellular function, leading to cyst formation and kidney enlargement.

While *PKD1* and *PKD2* account for the majority of ADPKD cases, variants in other genes have also been implicated in a small subset of patients. Although these genes are less frequent causes, they contribute to the clinical variability of ADPKD and broaden the spectrum of associated phenotypes [27]. However, due to the low prevalence of ADPKD in the UK Biobank dataset, only 154 patients were coded as either Q61.2 or Q61.3, traditional rare variant association tests can only identify *PKD1* and *PKD2* significantly after p-value adjustment. This limitation underscores the challenge of rare disease gene discovery and the potential value of leveraging phenotype-based similarity methods like PERADIGM to identify additional candidate genes associated with ADPKD-related phenotypes.

Before applying the PERADIGM approach, we first examined the intra-group similarity among individuals diagnosed with ADPKD and carriers of rare loss-of-function (LoF) variants in the *PKD1* and *PKD2* genes. Intra-group similarity is defined as the average pairwise phenotype similarity within a selected group, calculated based on each individual’s complete phenotype profile. We aimed to determine whether individuals within these two groups exhibited greater phenotypic similarity compared to randomly selected individuals. For the ADPKD group, the control group consisted of individuals without an ADPKD diagnosis in the inpatient EHR. For the variant carrier analysis, the control group included individuals without LoF or deleterious missense variants in the target gene.

We first considered ADPKD patients. As shown in fig 2a, there is an apparent separation of pairwise similarity scores between the ADPKD disease group and the ADPKD non-disease group. The ADPKD patients generally had higher pairwise similarity scores than non-patients, supported by the mean difference. This indicates that, on the phenotype level, ADPKD patients were more similar to each other compared to non-ADPKD individuals. These results suggest that the ADPKD patients shared some common and unique phenotypes which are significantly different from the non-patient group. We next considered the rare LoF variant carriers of *PKD1* and *PKD2*. As shown in fig 2b and fig 2c, *PKD1* and *PKD2* rare LoF variant carriers groups also exhibited more similar phenotypes. *PKD1* rare LoF variant carriers had a wider range of pairwise similarity scores, while the non-carrier group was more concentrated around 0.1. *PKD2* rare LoF variant carriers’ pairwise similarity score distribution showed a similar trend to ADPKD patients, compared to the controls, although the difference was more mitigated than in the ADPKD patients group. The *PKD1* and *PKD2* results show that while the pairwise similarity trends differed between the two genes’ rare LoF carriers, both had apparent differences compared to the control group. Moreover, the overall difference was less pronounced than the ADPKD patients group, suggesting that *PKD1* and *PKD2* only account for part of the phenotype patterns for ADPKD, and some other phenotype patterns cannot be explained by these two genes alone.

**Fig. 1:**
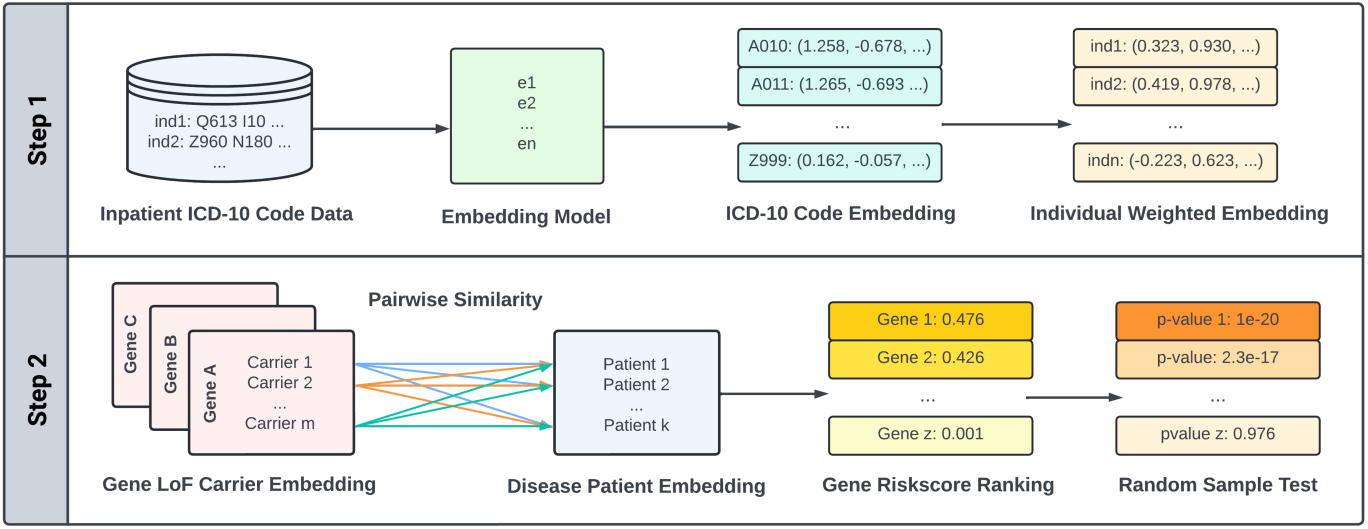
Overview of the PERADIGM framework. In Step 1, individual-level ICD-10 codes from EHR data are embedded using a Word2Vec model, and patient embeddings are generated as a weighted average of code embeddings, incorporating both disease relevance and phenotype prevalence. In Step 2, pairwise cosine similarity is computed between rare LoF variant carriers and disease patients. A gene-specific risk score is calculated based on the average similarity, and statistical significance is assessed via random sampling. This approach enables the identification of genes associated with disease-specific phenotypes.

**Fig. 2:**
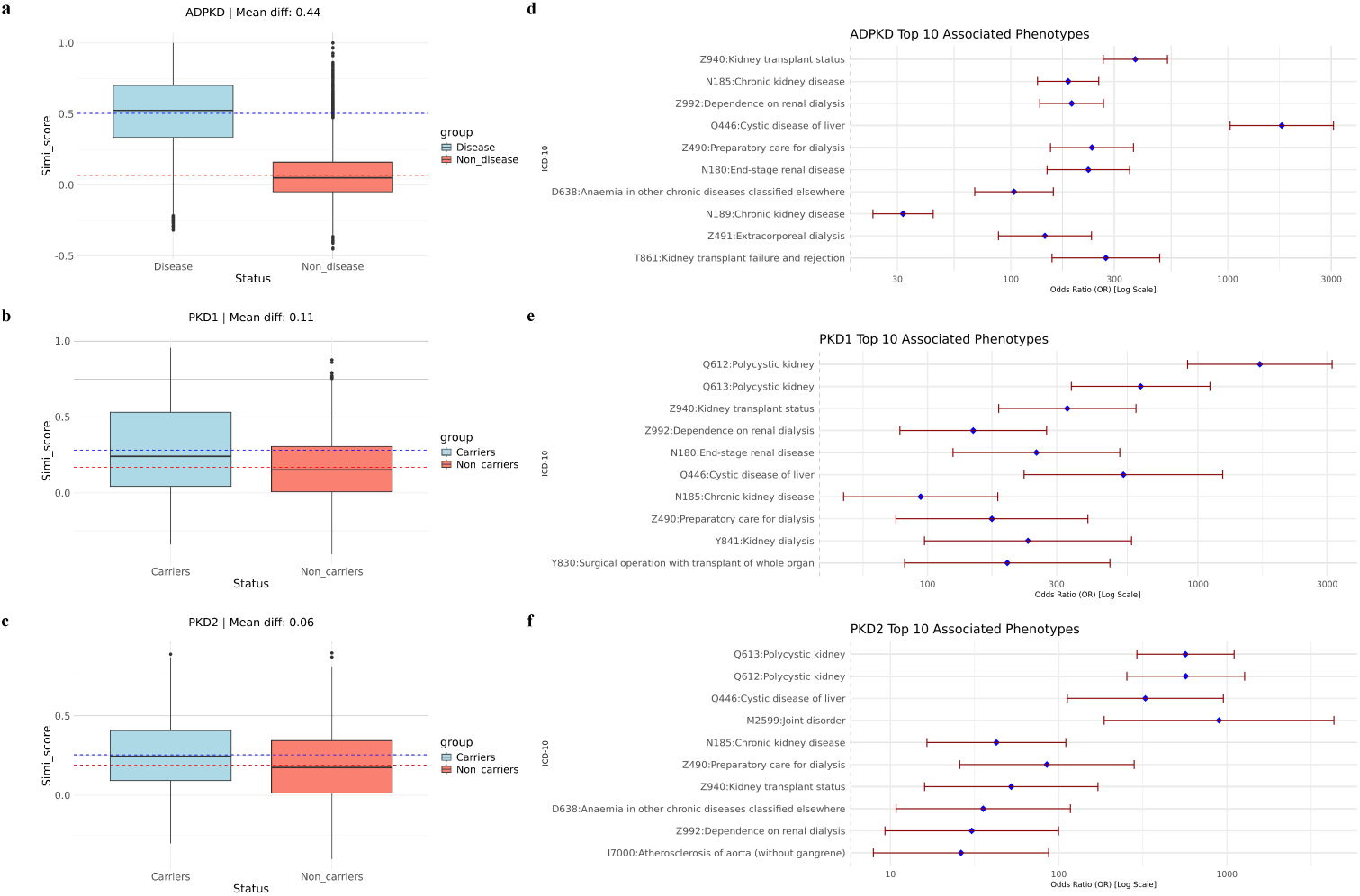
Intra-group similarity and significant phenotypes in the ADPKD analysis. **a.** Pairwise similarity scores within the ADPKD patient group and the control group. **b.** Pairwise similarity scores within the *PKD1* rare LoF variant carrier group and the control group. **c.** Pairwise similarity scores within the *PKD2* rare LoF variant carrier group and the control group. **d.** Top 10 significantly associated ICD-10 codes among ADPKD patients. **e.** Top 10 significantly associated ICD-10 codes among *PKD1* rare LoF variant carriers. **f.** Top 10 significantly associated ICD-10 codes among *PKD2* rare LoF variant carriers.

After analyzing the pairwise phenotype similarity among ADPKD patients, *PKD1* and *PKD2* genes, we next investigated which phenotypes contributed to the difference in phenotype patterns. In this analysis, we used logistic regression, adjusted for age and sex, to scan all the phenotypes available in the UK Biobank 200K dataset to identify phenotypes that are significantly associated with ADPKD. We considered 10,487 unique ICD-10 codes available in the UKBB 200K dataset.

For ADPKD, we identified 255 significantly associated phenotypes after *p*-value adjustment, excluding those present in only one patient. As illustrated in Fig. 2d, the top 10 phenotypes demonstrate strong associations with kidney diseases, cystic conditions, and common ADPKD complications [26]. These results show that, as expected, ADPKD patients not only share the primary diagnosis but also exhibit a constellation of related phenotypes. Among the significant associations, conditions such as hyperkalemia and urinary tract infections, both commonly observed in chronic kidney disease and ADPKD, were also identified (see Supplementary Materials). The ability of our approach to capture these well-known phenotypic associations suggests that such phenotype data could be leveraged to enhance individual-level phenotype embeddings.

We further investigated the phenotypes associated with *PKD1* and *PKD2* rare LoF carrier status. As shown in Fig. 2e and 2f, Q61.2 and Q61.3 were the two most significant phenotypes, as expected. Additionally, most of the top 10 significant phenotypes for *PKD1* and *PKD2* overlapped with those observed for ADPKD patients [8], further validating known genotype-phenotype relationships. Furthermore, we identified 68 significant phenotypes for *PKD1* and 15 for *PKD2* after p-value adjustment, which is substantially fewer than the 255 significant phenotypes identified for ADPKD patients. This discrepancy likely reflects the broader phenotypic spectrum captured in diagnosed ADPKD cases rather than solely the contributions of *PKD1* and *PKD2*.

Additionally, differences in coding practices among healthcare providers may contribute to variations in phenotype capture. Some providers may consistently document chronic kidney disease (CKD) as a general diagnosis during each visit while omitting more specific codes such as Q61.2 for ADPKD and other related findings. This limitation of ICD coding suggests that phenotype-based methods may help mitigate the effects of inconsistent coding practices. These results emphasize the potential of phenotype similarity-based approaches to improve the representation of individual disease profiles and enhance rare disease gene discovery.

After examining these phenotype patterns, we applied PERADIGM to all available genes in the UK Biobank 200K dataset to identify significant risk genes for ADPKD. After filtering out genes without rare LoF variant carriers, 17,792 genes remained. In our final analysis, we included only genes with at least ten rare LoF carriers, resulting in a total of 14,929 genes.

As shown in Fig. 3a and Table 1, genome-wide scanning with PERADIGM identified six significant genes associated with ADPKD-related phenotypes after p-value adjustment. In comparison, SKAT-O identified only *PKD1* and *PKD2* after p-value adjustment (Fig. 3b), whereas PERADIGM identified four additional candidate genes.

**Table 1:** Summary of significant genes associated with ADPKD-specific phenotypes identified by PERADIGM. Each row represents a gene identified by PERADIGM. The columns show the gene name, the number of individuals carrying rare LoF variants in that gene (Carrier Number), the number of these carriers who were also diagnosed with ADPKD (Overlap with Disease), the computed risk score, the raw p-value from the similarity-based test, and the adjusted p-value after multiple testing correction.

| Gene | Carrier Number | Overlap with Disease | Riskscore | Raw P-value | Adjusted P-value |
| --- | --- | --- | --- | --- | --- |
| *PKD1* | 43 | 17 | 0.31 | $<1.0 \times 10^{-25}$ | $<1.0 \times 10^{-25}$ |
| *PKD2* | 33 | 9 | 0.22 | $1.81 \times 10^{-14}$ | $2.61 \times 10^{-10}$ |
| *TET2* | 186 | 1 | 0.11 | $2.11 \times 10^{-11}$ | $3.04 \times 10^{-7}$ |
| *ASXL1* | 181 | 1 | 0.11 | $5.44 \times 10^{-9}$ | $7.83 \times 10^{-5}$ |
| *CGNL1* | 54 | 1 | 0.13 | $2.07 \times 10^{-6}$ | $2.99 \times 10^{-2}$ |
| *IFT140* | 349 | 7 | 0.08 | $3.36 \times 10^{-6}$ | $4.84 \times 10^{-2}$ |

**Fig. 3:**
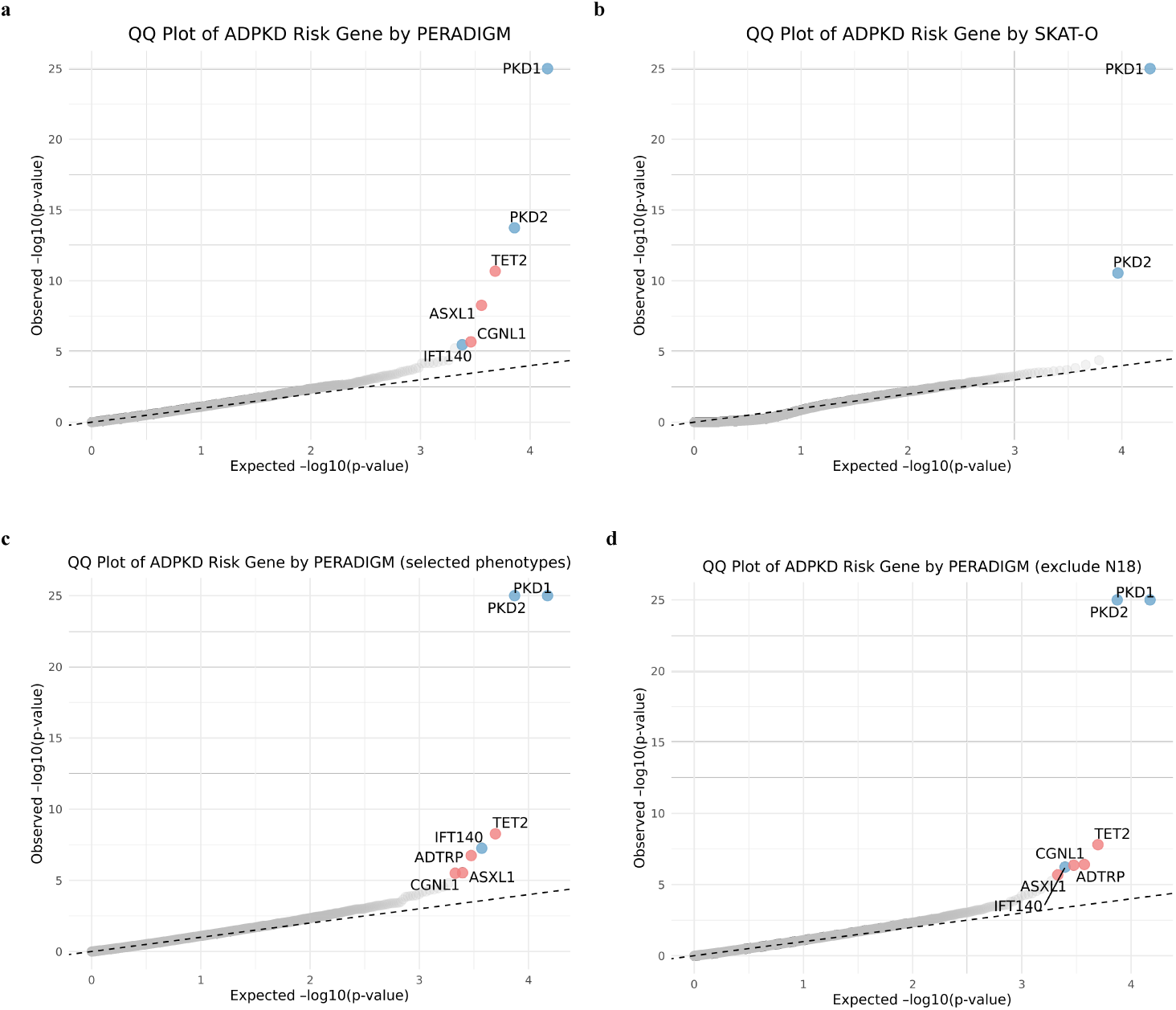
QQ plots of ADPKD analysis. **a.** QQ plot of ADPKD risk genes identified by PERADIGM. **b.** QQ plot of ADPKD risk genes identified by SKAT-O. **c.** QQ plot from PERADIGM using pre-selected ADPKD-related phenotypes for embedding the patient group. **d.** QQ plot from PERADIGM after excluding CKD patients from the analysis. Blue points indicate significant genes that were also verified in previous studies, while red points represent newly identified significant genes.

Among these genes, *IFT140* is a well-established gene in ciliopathies that has recently been implicated as a monogenic cause of renal cyst formation [28]. Mutations in *IFT140* can disrupt cilia structure and function. However, SKAT-O failed to detect *IFT140* as significant (raw p-value: 0.00146, adjusted p-value: 0.58, rank: 46), likely due to the small number of *IFT140* rare LoF variant carriers among individuals with ADPKD diagnosed (only seven individuals). This limited overlap reduces statistical power in binary status-based association tests like SKAT-O. In contrast, PERADIGM integrates phenotype information across all ADPKD-related phenotypes for each individual with rare LoF variants, providing a more comprehensive assessment beyond binary disease status. As for *TET2*, although it is not a known ADPKD gene, it has been implicated in kidney disease, particularly in acute kidney injury (AKI) and chronic kidney disease, both of which are commonly observed in individuals with ADPKD-related phenotypes [29, 30]. *ASXL1*, another significant gene, interacts with

WTIP, a binding protein critical for kidney podocyte development. Disruptions in *ASXL1* can result in smaller kidneys and reduced glomerular podocyte volume [31]. *CGNL1* was identified as a candidate gene for kidney injury and hypertension, with studies showing that *CGNL1* knockout can reduce renal injury and salt-sensitive blood pressure [32]. To further compare PERADIGM with SKAT-O, we examined overlapping genes among the top 100, 200, and 500 genes ranked by raw p-values. The results showed 6, 8, and 26 overlapping genes, respectively. The relatively small number of shared genes suggests that PERADIGM prioritizes different aspects of gene significance compared to SKAT-O, potentially capturing distinct biological signals related to ADPKD-specific phenotypes. These results demonstrate that incorporating phenotype similarity into gene discovery can enhance the identification of candidate genes associated with ADPKD-related phenotypes. By leveraging phenotype embeddings, PERADIGM not only validates known genetic contributors but also uncovers additional genes that may influence specific clinical manifestations, providing deeper insights into the phenotypic complexity of ADPKD.

Beyond the significant genes identified by PERADIGM (Table 1), additional evidence supports its enhanced power. Several genes previously linked to ADPKD, including *ALG8* [33], *ALG9* [34], *COL4A1* [35], and *SEC63* [36], showed marginally significant associations when analyzed with PERADIGM. These genes had minimal overlap between rare LoF variant carriers and ADPKD diagnosis (*ALG8* : 1, *ALG9* : 0, *COL4A1* : 0, *SEC63* : 2), which likely contributed to their lack of significance in SKAT-O. Among them, only *SEC63* achieved marginal significance in SKAT-O, whereas PERADIGM identified marginal associations for all four genes (Table 2). This result highlights how limited overlap between rare variant carriers and ADPKD diagnoses can reduce SKAT-O’s power; however, this may reflect incomplete ICD coding, undiagnosed cases, or reduced penetrance rather than a true absence of association. In contrast, PERADIGM remains effective by leveraging broader phenotype patterns.

**Table 2:** Marginal p-values of ADPKD for PERADIGM and SKAT-O. Each row corresponds to a gene. The columns show the gene name, the number of variant carriers (Carrier Number), the number of carriers overlapping with ADPKD diagnoses (Overlap with Disease), the p-value from PERADIGM, and the p-value from SKAT-O.

| Gene | Carrier Number | Overlap with Disease | P-value by PERADIGM | P-value by SKAT-O |
| --- | --- | --- | --- | --- |
| *ALG8* | 149 | 1 | $1.57 \times 10^{-2}$ | $4.75 \times 10^{-1}$ |
| *ALG9* | 32 | 0 | $4.10 \times 10^{-2}$ | $6.16 \times 10^{-1}$ |
| *COL4A1* | 20 | 0 | $3.69 \times 10^{-3}$ | $1.00 \times 10^0$ |
| *SEC63* | 46 | 2 | $9.71 \times 10^{-4}$ | $1.29 \times 10^{-3}$ |

Moreover, we conducted an additional analysis to refine the ADPKD-related phenotype representation for diagnosed patients. Instead of using the entire phenotype profile, we preselected ADPKD-related phenotypes based on prior expertise when generating patient group embeddings. This stricter phenotype selection enhances specificity, further validates PERADIGM’s results, and provides more explainable associations between significant genes and ADPKD-specific phenotypes. For ADPKD, we selected ADPKD, aneurysms, and liver cysts as key ADPKD-related phenotypes. As shown in Fig. 3c and Table 3, the significant genes identified remained nearly the same as those found using PERADIGM without phenotype preselection, with one additional identified gene *ADTRP*. This consistency indicates that both approaches yield robust gene discovery results and highlights PERADIGM’s ability to automatically assign higher weights to phenotypes most relevant to ADPKD. However, the results using the preselected phenotype group showed a stronger association of *PKD1* and *PKD2* with ADPKD-related phenotypes (both adjusted p-values = 0.00), and the ranking of the seven significant genes changed, with *IFT140* becoming more significant. This shift suggests that preselecting ADPKD-related phenotypes can slightly enhance statistical power, but the overall results remain consistent with those obtained without phenotype preselection, demonstrating the robustness of PERADIGM.

**Table 3:** Risk scores and p-values of significant genes for ADPKD based on comparison group with selected phenotypes. Each row represents a gene. Columns show the gene name, the number of variant carriers (Carrier Number), the number of carriers overlapping with ADPKD diagnosis (Overlap with Disease), the calculated risk score, the raw p-value, and the adjusted p-value after multiple testing corrections.

| Gene | Carrier Number | Overlap with Disease | Riskscore | Raw P-value | Adjusted P-value |
| --- | --- | --- | --- | --- | --- |
| *PKD1* | 43 | 17 | 0.36 | $<1.0 \times 10^{-25}$ | $<1.0 \times 10^{-25}$ |
| *PKD2* | 33 | 9 | 0.25 | $<1.0 \times 10^{-25}$ | $<1.0 \times 10^{-25}$ |
| *TET2* | 186 | 1 | 0.10 | $5.38 \times 10^{-9}$ | $8.03 \times 10^{-5}$ |
| *IFT140* | 349 | 7 | 0.08 | $5.57 \times 10^{-8}$ | $8.32 \times 10^{-4}$ |
| *ADTRP* | 27 | 0 | 0.18 | $1.82 \times 10^{-7}$ | $2.71 \times 10^{-3}$ |
| *ASXL1* | 181 | 1 | 0.09 | $2.94 \times 10^{-6}$ | $4.39 \times 10^{-2}$ |
| *CGNL1* | 54 | 1 | 0.13 | $3.18 \times 10^{-6}$ | $4.75 \times 10^{-2}$ |

In addition, ADPKD is a specific subtype of chronic kidney disease (CKD), and the two conditions share many overlapping phenotypes. While CKD is often present in ADPKD patients, many of the complications and phenotypes observed in CKD can arise from a variety of non-genetic causes. To distinguish genes associated specifically with ADPKD-related phenotypes from those potentially confounded by common CKD, we conducted an additional analysis. Specifically, we excluded individuals diagnosed with CKD (ICD-10 code: N18) who did not also have a diagnosis of ADPKD. This filtering step aimed to remove individuals with non-ADPKD forms of CKD that could obscure the genetic signals specific to ADPKD. We then applied PERADIGM using this refined comparison group to identify genes associated with ADPKD-specific phenotypes. As shown in Fig. 3d and Table 4, all previously identified genes remained significant, consistent with our original findings. This result demonstrates the robustness of PERADIGM and its ability to detect ADPKD-related genetic associations even after removing potentially confounding CKD cases.

**Table 4:** Risk scores and p-values of significant genes for ADPKD based on comparison group excluding CKD patients. Each row represents a gene. Columns show the gene name, the number of variant carriers (Carrier Number), the number of carriers overlapping with ADPKD diagnosis (Overlap with Disease), the calculated risk score, the raw p-value, and the adjusted p-value after multiple testing corrections.

| Gene | Carrier Number | Overlap with Disease | Riskscore | Raw P-value | Adjusted P-value |
| --- | --- | --- | --- | --- | --- |
| *PKD1* | 42 | 17 | 0.35 | $<1.0 \times 10^{-25}$ | $<1.0 \times 10^{-25}$ |
| *PKD2* | 33 | 9 | 0.24 | $<1.0 \times 10^{-25}$ | $<1.0 \times 10^{-25}$ |
| *TET2* | 182 | 1 | 0.10 | $1.57 \times 10^{-8}$ | $2.35 \times 10^{-4}$ |
| *ADTRP* | 26 | 0 | 0.17 | $3.89 \times 10^{-7}$ | $5.81 \times 10^{-3}$ |
| *CGNL1* | 54 | 1 | 0.13 | $4.35 \times 10^{-7}$ | $6.49 \times 10^{-3}$ |
| *IFT140* | 343 | 7 | 0.07 | $5.86 \times 10^{-7}$ | $8.75 \times 10^{-3}$ |
| *ASXL1* | 178 | 1 | 0.09 | $2.09 \times 10^{-6}$ | $3.11 \times 10^{-2}$ |

### 2.3 Marfan syndrome

We next applied PERADIGM to Marfan syndrome [37], an autosomal dominant connective tissue disorder primarily caused by mutations in *FBN1*. This gene encodes fibrillin-1, a crucial protein for elastic fiber formation. Defective fibrillin-1 disrupts connective tissue integrity, leading to characteristic features such as tall stature, long limbs, flexible joints, lens dislocation, and serious cardiovascular complications, including aortic aneurysms and dissections. While *FBN1* mutations are the main cause, other genes have been implicated in Marfan syndrome and Marfan-like conditions, including *TGFBR1* and *TGFBR2* [38]. Mutations in these genes can also impair connective tissue function, contributing to the syndrome’s clinical manifestations. This suggests that beyond *FBN1*, multiple genes may influence connective tissue integrity and related pathways. In the UK Biobank 200K dataset, rare variant association tests such as SKAT-O, SKAT, and Burden tests identified only *FBN1* as a significant gene after p-value adjustment. We applied PERADIGM to Marfan syndrome to explore additional candidate genes associated with Marfan-related phenotypes.

We first examined the intra-group similarity among Marfan syndrome patients and rare LoF variant carriers of *FBN1*. In the UK Biobank dataset, 42 individuals were diagnosed with Marfan syndrome (ICD-10 code: Q87.4). As shown in Fig. 4a, the intra-group similarity scores of Marfan syndrome patients differed substantially from those of non-patients. The boxplot revealed an average pairwise similarity score difference of 0.55, which is notably higher than that for ADPKD. This strong intra-group similarity indicates that Marfan syndrome patients exhibit distinct and characteristic phenotype patterns. We then analyzed the intra-group similarity scores of 11 individuals carrying rare LoF variants in *FBN1*, five of whom were diagnosed with Marfan syndrome. Although the number of carriers was small, Fig. 4b shows a clear difference between carriers and controls. The similarity scores were sparse due to the limited number of carriers, but the overall trend suggests slightly higher intra-group similarity among *FBN1* carriers. Notably, the phenotype pattern of *FBN1* rare LoF carriers was weaker than that of diagnosed Marfan syndrome patients, as reflected by lower intra-group similarity scores. This suggests that *FBN1* exhibits incomplete penetrance for Marfan syndrome and that additional phenotype patterns may exist beyond those explained by *FBN1* alone. These findings indicate the potential involvement of other genetic contributors in Marfan syndrome.

**Fig. 4:**
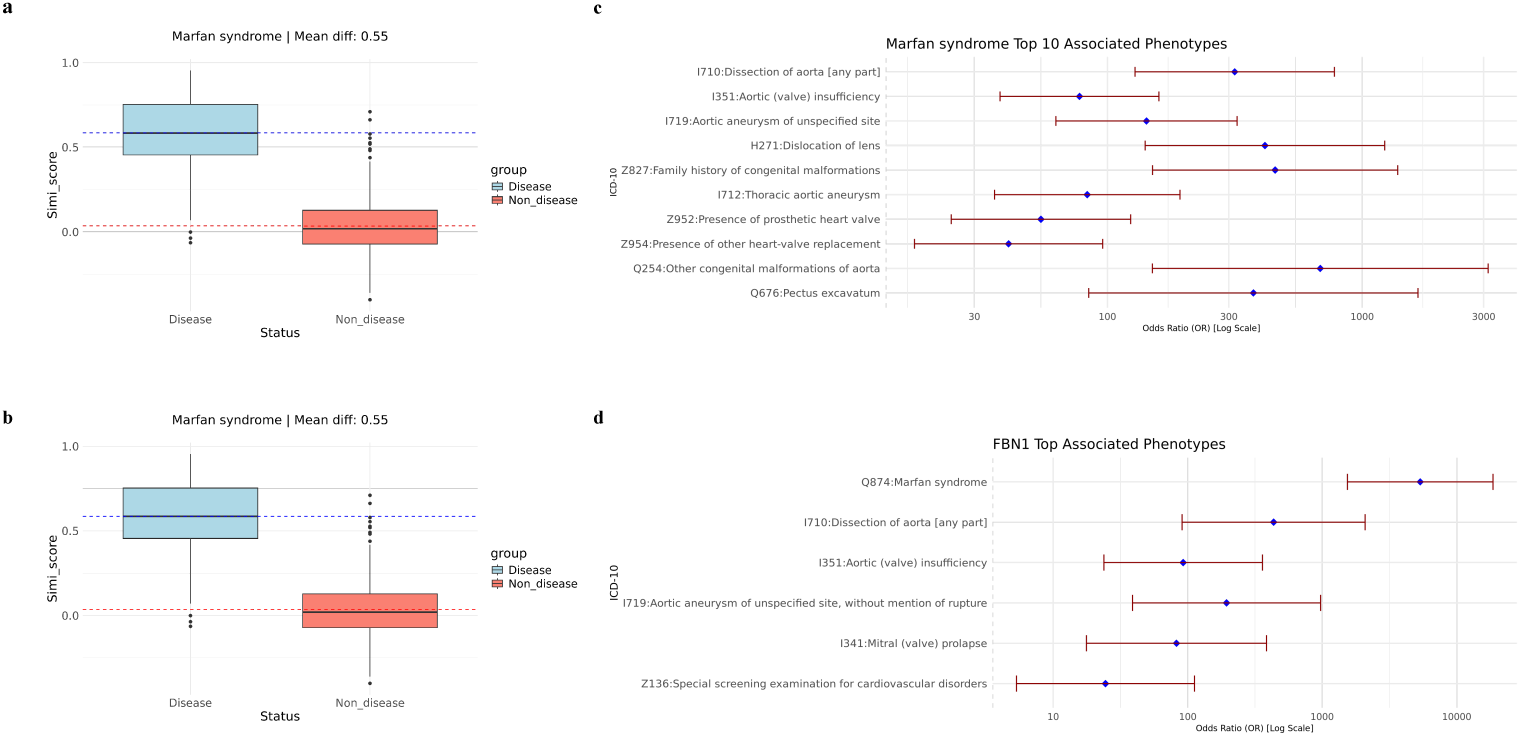
Intra-group similarity and significant phenotypes in the Marfan syndrome analysis. **a.** Pairwise similarity scores within the Marfan syndrome patient group and the control group. **b.** Pairwise similarity scores within the *FBN1* variantcarrier group and the control group. **c.** Top 10 significantly associated ICD-10 codes among Marfan syndrome patients. **d.** Top 10 significantly associated ICD-10 codes among *FBN1* variant carriers.

Next, we examined the significantly associated phenotypes of Marfan syndrome and *FBN1* rare LoF variant carriers to better understand the phenotypic contributors to Marfan syndrome. After p-value adjustment, 51 significantly associated phenotypes remained, excluding those present in only one patient. Fig. 4c illustrates the top 10 significant phenotypes associated with Marfan syndrome, which are predominantly related to the aorta, congenital heart disease, and blood abnormalities—all common complications of the disorder [39]. Additionally, some significant phenotypes without a direct known relationship to Marfan syndrome were identified, such as retinal detachment with retinal break and ingrowing nails (see Supplementary Materials). This pattern of significant phenotypes, comprising both common complications and less typical associations, is consistent with our findings in ADPKD. We then investigated whether *FBN1* rare LoF carriers exhibited similar significant phenotypes to Marfan syndrome patients. Due to the limited number of carriers, only six phenotypes were significantly associated with *FBN1* rare LoF variant status. Despite the small sample size, all six phenotypes were known complications of Marfan syndrome (Fig. 4d) and were also observed in diagnosed patients. However, many additional significant phenotypes observed in Marfan syndrome patients were absent in *FBN1* rare LoF carriers. This indicates that *FBN1* exhibits incomplete penetrance for Marfan syndrome, suggesting that other factors may influence the broader spectrum of significant phenotypes associated with the disease.

After exploring the phenotype patterns of Marfan syndrome and *FBN1* rare LoF carriers, we applied PERADIGM to scan all genes in the UK Biobank 200K dataset to identify significant genes associated with Marfan syndrome-specific phenotypes [40]. As shown in Fig. 5a and Table 5, the whole-genome scan identified four significant genes after p-value adjustment, whereas SKAT-O identified only *FBN1* (Fig. 5b). While *FBN1* is a well-established gene for Marfan syndrome, the other three significant genes were newly identified by our method. These novel findings suggest that additional genetic contributors may be involved in Marfan syndrome-specific phenotypes, expanding the understanding of its genetic landscape. These results demonstrate that PERADIGM can effectively identify candidate genes for rare disease-related phenotypes by integrating comprehensive phenotype information.

**Fig. 5:**
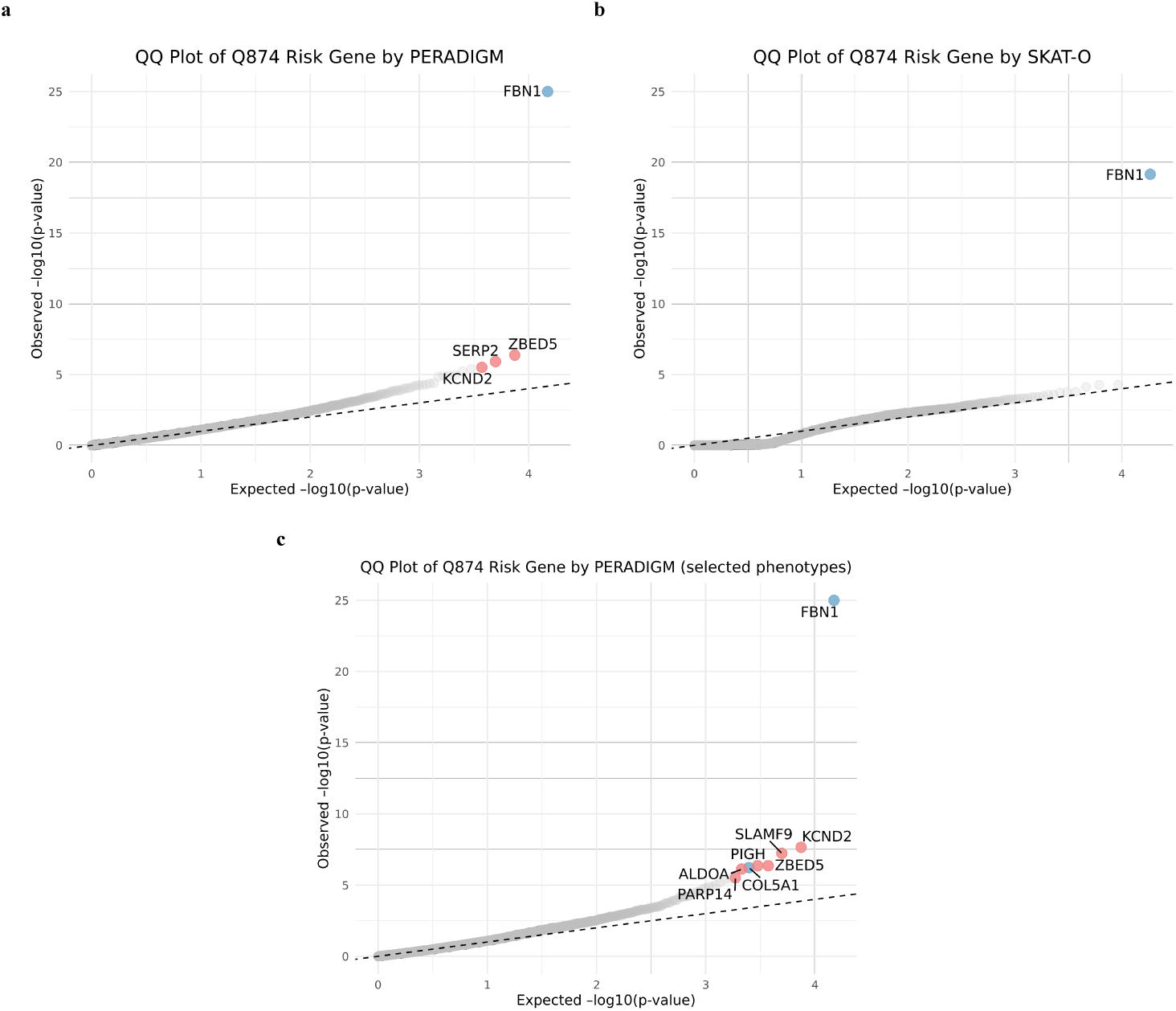
QQ plots of Marfan syndrome analysis. **a.** QQ plot of Marfan syndrome risk genes identified by PERADIGM. **b.** QQ plot of Marfan syndrome risk genes identified by SKAT-O. **c.** QQ plot from PERADIGM using pre-selected Marfan syndrome-related phenotypes for embedding the patient group. Blue points indicate significant genes that were also verified in previous studies, while red points represent newly identified significant genes.

**Table 5:** Summary of significant genes associated with Marfan syndrome-specific phenotypes identified by PERADIGM. Each row represents a gene identified by PERADIGM. The columns show the gene name, the number of individuals carrying variants in that gene (Carrier Number), the number of these carriers who were also diagnosed with Marfan syndrome (Overlap with Disease), the computed risk score, the raw p-value from the similarity-based test, and the adjusted p-value after multiple testing correction.

| Gene | Carrier Number | Overlap with Disease | Riskscore | Raw P-value | Adjusted P-value |
| --- | --- | --- | --- | --- | --- |
| *FBN1* | 11 | 5 | 0.33 | $<1.0 \times 10^{-25}$ | $<1.0 \times 10^{-25}$ |
| *ZBED5* | 53 | 0 | 0.10 | $4.21 \times 10^{-7}$ | $6.28 \times 10^{-3}$ |
| *SERP2* | 11 | 0 | 0.17 | $1.22 \times 10^{-6}$ | $1.82 \times 10^{-2}$ |
| *KCND2* | 12 | 0 | 0.16 | $3.08 \times 10^{-6}$ | $4.60 \times 10^{-2}$ |

We also conducted an additional analysis to refine the representation of the Marfan syndrome phenotype in diagnosed patients. Instead of using the entire phenotype profile, we embedded the diagnosed patients using only their longitudinal ICD-10 codes for Marfan syndrome, ensuring that the embeddings were derived solely from repeated Marfan syndrome diagnoses over time. This stricter embedding process focuses more specifically on Marfan syndrome itself, providing more explainable results. As shown in Fig. 5c and Table 6, eight genes were significantly associated with Marfan syndrome-specific phenotypes. These include the three previously identified genes, indicating that refining phenotype representation can help uncover additional relevant genes.

**Table 6:** Risk scores and p-values of significant genes for Marfan syndrome based on comparison group with selected phenotypes. Each row represents a gene. Columns show the gene name, the number of variant carriers (Carrier Number), the number of carriers overlapping with Marfan syndrome diagnosis (Overlap with Disease), the calculated risk score, the raw p-value, and the adjusted p-value after multiple testing corrections.

| Gene | Carrier Number | Overlap with Disease | Riskscore | Raw P-value | Adjusted P-value |
| --- | --- | --- | --- | --- | --- |
| *FBN1* | 11 | 5 | 0.38 | $<1.0 \times 10^{-25}$ | $<1.0 \times 10^{-25}$ |
| *KCND2* | 12 | 0 | 0.18 | $2.22 \times 10^{-8}$ | $3.32 \times 10^{-4}$ |
| *SLAMF9* | 43 | 1 | 0.09 | $5.91 \times 10^{-8}$ | $8.82 \times 10^{-4}$ |
| *ZBED5* | 53 | 0 | 0.07 | $4.34 \times 10^{-7}$ | $6.47 \times 10^{-3}$ |
| *PIGH* | 39 | 0 | 0.09 | $4.35 \times 10^{-7}$ | $6.50 \times 10^{-3}$ |
| *COL5A1* | 26 | 0 | 0.11 | $5.93 \times 10^{-7}$ | $8.85 \times 10^{-3}$ |
| *ALDOA* | 17 | 0 | 0.13 | $7.48 \times 10^{-7}$ | $1.12 \times 10^{-2}$ |
| *PARP14* | 199 | 0 | 0.03 | $3.08 \times 10^{-6}$ | $4.60 \times 10^{-2}$ |

Among the four newly identified genes, *COL5A1*, primarily associated with Ehlers-Danlos syndrome (EDS), has been reported in cases where EDS and Marfan syndrome are difficult to distinguish due to overlapping connective tissue abnormalities [41]. *COL5A1* mutations result in abnormal type V collagen production, leading to symptoms that resemble certain aspects of Marfan syndrome. In our dataset, none of the 26 *COL5A1* rare LoF carriers overlapped with diagnosed Marfan syndrome patients, highlighting the limitations of traditional rare variant association methods in detecting such associations. The ICD-10 coding system sometimes struggles to distinguish rare diseases like Marfan syndrome and EDS, leading to potential misdiagnoses. By leveraging full phenotype embeddings, our approach can help mitigate these issues by capturing broader phenotype patterns. These results further demonstrate that PERADIGM can effectively identify genetic contributors to Marfan syndrome-specific phenotypes by integrating comprehensive phenotype information. By utilizing phenotype embeddings, PERADIGM enables the discovery of genes that contribute to disease manifestations through shared pathways or overlapping syndromes, even when there is no direct overlap between gene carriers and diagnosed patients.

Additionally, *TGFBR2* is associated with Loeys-Dietz syndrome, a condition that shares several features with Marfan syndrome, including aortic aneurysms and skeletal abnormalities [42]. In our analysis, *TGFBR2* ranked 16th in the whole-genome scan (raw p-value = 6.1 → 10^→5^), whereas SKAT-O ranked 7571st (raw p-value = 9.3 → 10^→1^). Mutations in *TGFBR2* disrupt the TGF-*ω* signaling pathway, which is essential for connective tissue maintenance and repair, resulting in clinical features that overlap with Marfan syndrome. In our UK Biobank dataset, none of the 12 *TGFBR2* rare LoF carriers were diagnosed with Marfan syndrome, preventing traditional binary-based methods from detecting this association. However, PERADIGM ranked *TGFBR2* among the top genes and achieved marginal significance, suggesting its potential involvement in Marfan syndrome-specific phenotypes despite the lack of direct overlap between gene carriers and diagnosed patients.

### 2.4 NF1

Neurofibromatosis type 1 (NF1) [43] is a complex genetic disorder characterized by the formation of multiple benign tumors, primarily neurofibromas, along nerves in the skin, brain, and other tissues. The disorder is primarily caused by mutations in the *NF1* gene, which encodes neurofibromin, a tumor suppressor protein that regulates cell growth and division. While *NF1* mutations are the main genetic driver, recent studies suggest that additional genes may influence disease severity and phenotypic variability among NF1 patients. Identifying such modifier genes could improve our understanding of NF1 pathogenesis and facilitate more personalized diagnostic and therapeutic strategies. In the UK Biobank 200K European ancestry dataset, there are 63 NF1 cases, limiting the statistical power of traditional rare variant association tests to detect genetic factors beyond *NF1* itself. To address this limitation, we applied PERADIGM to identify genetic contributors that may influence NF1-specific phenotypes or modify the clinical presentation, expanding beyond the established role of *NF1*.

We first examined the intra-group similarity among NF1 patients and rare LoF carriers of the *NF1* gene. The UK Biobank 200K European ancestry dataset contained 63 individuals diagnosed with NF1 (ICD-10 code: Q85.0). As shown in Fig. 6a, NF1 patients exhibited significantly higher intra-group similarity compared to the control group, indicating distinct phenotypic characteristics. We then analyzed intra-group similarity for 114 individuals carrying rare LoF variants in *NF1*, among whom only 19 had a documented NF1 diagnosis. This suggests that while *NF1* mutations are the primary cause of NF1, many *NF1* rare LoF variant carriers exhibit reduced or incomplete penetrance and conversely, some diagnosed patients do not carry detectable *NF1* rare LoF variants. As depicted in Fig. 6b, the difference in intra-group similarity between *NF1* rare LoF carriers and controls was less pronounced than that observed for diagnosed NF1 patients. These findings are consistent with results from ADPKD and Marfan syndrome, suggesting that additional genetic or environmental factors likely influence NF1 disease manifestation beyond *NF1* mutations alone.

**Fig. 6:**
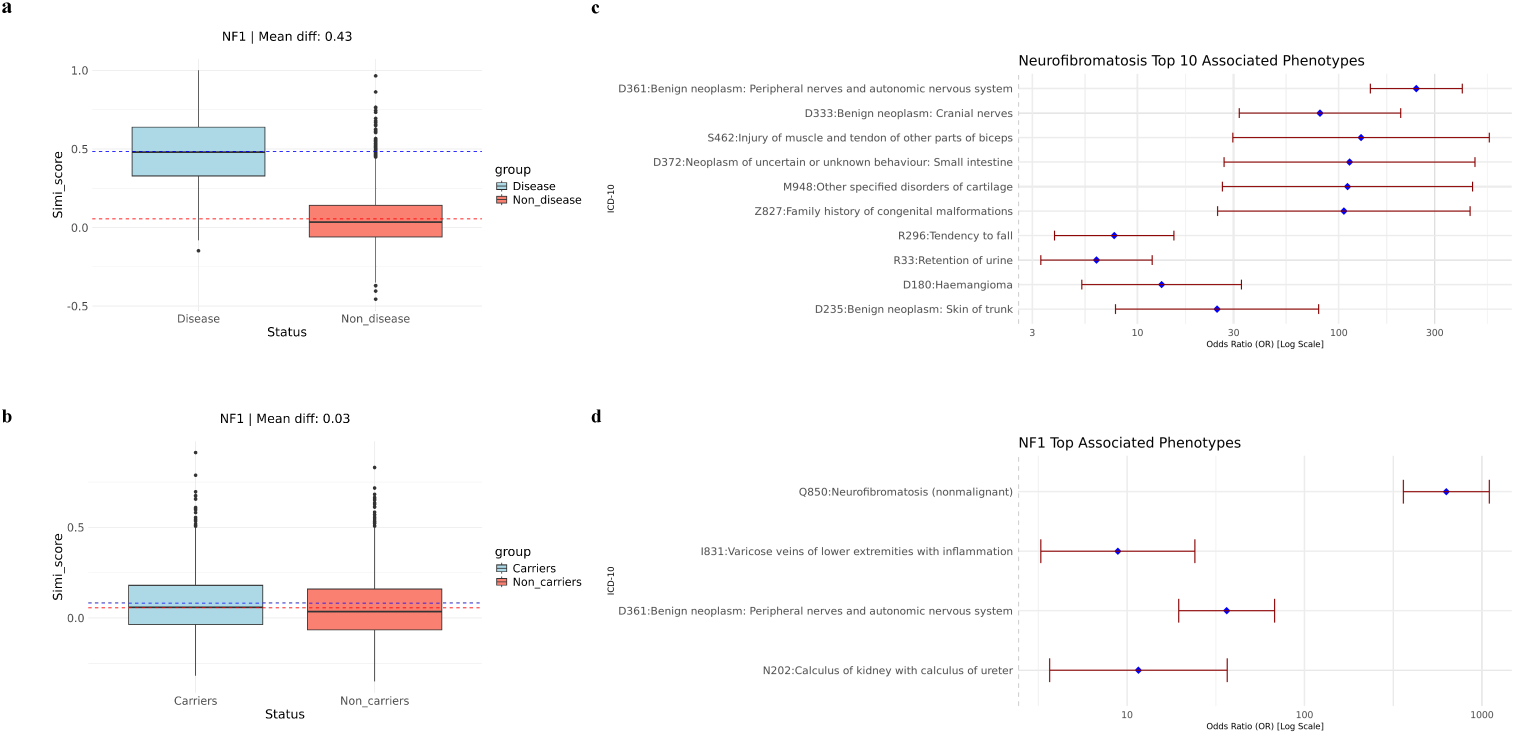
Intra-group similarity and significant phenotypes in the NF1 analysis. **a.** Pairwise similarity scores within the NF1 patient group and the control group. **b.** Pairwise similarity scores within the *NF1* variant carrier group and the control group. **c.** Top 10 significantly associated ICD-10 codes among NF1 patients. **d.** Top significantly associated ICD-10 codes among *NF1* variant carriers.

We examined phenotypes significantly associated with NF1 disease and *NF1* rare LoF variant carrier status, identifying 38 phenotypes associated with NF1 disease(Fig. 6c). The top 10 significant phenotypes closely aligned with known clinical manifestations, reinforcing the multisystem involvement characteristic of NF1 [44]. These phenotypes primarily affected the nervous system, skin, and skeleton, with some involvement of other organ systems. This comprehensive phenotypic profile enhances our understanding of NF1’s diverse clinical presentation. In contrast, despite a larger number of *NF1* rare LoF variant carriers (114) compared to diagnosed NF1 patients, only four phenotypes remained significant after p-value adjustment(Fig. 6d). As expected, NF1 disease itself was the most significant phenotype, followed by D36.1, which is directly related to NF1. Interestingly, the remaining two significant phenotypes are not commonly associated with NF1, suggesting potential novel genotype-phenotype correlations or incidental findings. NF1 patients exhibit a distinct and consistent phenotypic profile, whereas *NF1* rare LoF variant carriers present a more variable and diluted phenotype pattern. Apart from the strong association with NF1 disease itself, other phenotype relationships in carriers were less pronounced. This finding highlights the complexity of genotype-phenotype relationships in NF1 and suggests that additional genetic modifiers or environmental factors may influence disease manifestation.

Following our exploration of phenotype patterns in NF1 disease patients and *NF1* rare LoF variant carriers, we applied PERADIGM to scan all available genes in our dataset for associations with NF1-specific phenotypes. We then compared our results with those obtained using SKAT-O. As shown in Fig 7a, after p-value adjustment, PERADIGM identified only *NF1* as a statistically significant gene associated with NF1-specific phenotypes, consistent with SKAT-O results (Fig. 7b). Both methods converged on *NF1* as the sole significant genetic factor in our dataset. The difficulty in identifying additional significant genes may be due to the relatively diluted phenotypic patterns observed among *NF1* rare LoF variant carriers, as noted in our previous analyses. This suggests that while *NF1* is the primary genetic driver, the complexity of NF1 disease manifestation may involve additional genetic modifiers or environmental influences that remain undetected with current sample sizes and methods.

**Fig. 7:**
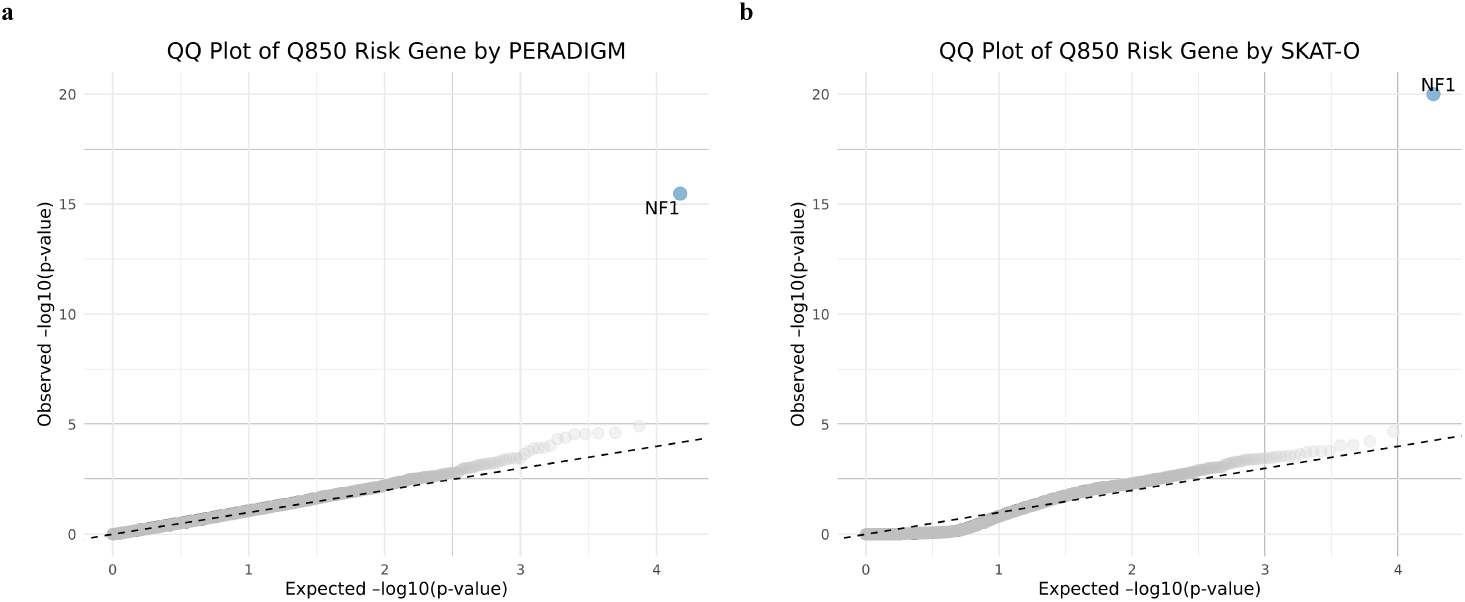
QQ plots of NF1 analysis. **a.** QQ plot of NF1 risk genes identified by PERADIGM. **b.** QQ plot of NF1 risk genes identified by SKAT-O.

From the QQ plot, we observed a departure from the central line at the tail using PERADIGM, suggesting potential additional signals. The top 10 genes are listed in Table 7. Among them, *P2RX4* encodes a purinergic receptor involved in cell signaling, particularly within the nervous system [45]. Impaired P2X receptor signaling has been reported in microglia with *NF1* mutations, highlighting the importance of purinergic receptors in *NF1* -related neurological abnormalities. Given its role, *P2RX4* may contribute to disrupted signaling pathways affecting microglial motility and phagocytosis in *NF1* [46]. Although these genes do not reach statistical significance after p-value adjustment, they may still provide valuable insights into genetic factors influencing *NF1* -specific phenotypes. Further investigation is needed to determine their potential contributions to *NF1* pathophysiology.

**Table 7:**
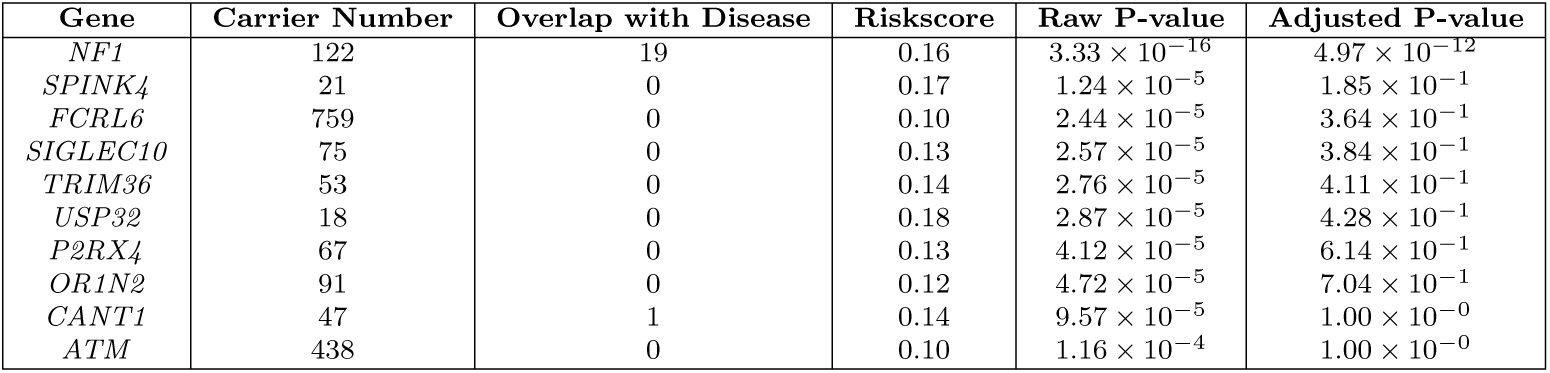
Risk scores and p-values of top 10 genes for NF1 disease. Each row represents a gene. Columns show the gene name, the number of variant carriers (Carrier Number), the number of carriers overlapping with NF1 diagnosis (Overlap with Disease), the calculated risk score, the raw p-value, and the adjusted p-value after multiple testing corrections.

| Gene | Carrier Number | Overlap with Disease | Riskscore | Raw P-value | Adjusted P-value |
| --- | --- | --- | --- | --- | --- |
| *NF1* | 122 | 19 | 0.16 | $3.33 \times 10^{-16}$ | $4.97 \times 10^{-12}$ |
| *SPINK4* | 21 | 0 | 0.17 | $1.24 \times 10^{-5}$ | $1.85 \times 10^{-1}$ |
| *FCRL6* | 759 | 0 | 0.10 | $2.44 \times 10^{-5}$ | $3.64 \times 10^{-1}$ |
| *SIGLEC10* | 75 | 0 | 0.13 | $2.57 \times 10^{-5}$ | $3.84 \times 10^{-1}$ |
| *TRIM36* | 53 | 0 | 0.14 | $2.76 \times 10^{-5}$ | $4.11 \times 10^{-1}$ |
| *USP32* | 18 | 0 | 0.18 | $2.87 \times 10^{-5}$ | $4.28 \times 10^{-1}$ |
| *P2RX4* | 67 | 0 | 0.13 | $4.12 \times 10^{-5}$ | $6.14 \times 10^{-1}$ |
| *OR1N2* | 91 | 0 | 0.12 | $4.72 \times 10^{-5}$ | $7.04 \times 10^{-1}$ |
| *CANT1* | 47 | 1 | 0.14 | $9.57 \times 10^{-5}$ | $1.00 \times 10^{-0}$ |
| *ATM* | 438 | 0 | 0.10 | $1.16 \times 10^{-4}$ | $1.00 \times 10^{-0}$ |

Furthermore, we examined *SPRED1* [47] and *LZTR1* [48], two genes previously implicated as potential contributors to NF1-specific phenotypes. Although neither gene reached statistical significance in PERADIGM or SKAT-O after p-value correction, our analysis revealed that *SPRED1* had 17 rare LoF variant carriers, none of whom were diagnosed with NF1 in our dataset. Similarly, *LZTR1* had 367 rare LoF variant carriers, with only one individual diagnosed with NF1. As shown in Table 8, both *SPRED1* and *LZTR1* exhibited marginal significance in PERADIGM, whereas SKAT-O identified only *LZTR1* as marginally significant. This discrepancy arises because SKAT-O relies heavily on direct overlap between LoF variant carriers and diagnosed patients, whereas PERADIGM can detect associations by leveraging shared phenotypic patterns. For example, despite the lack of overlap between *SPRED1* carriers and NF1 patients, PERADIGM still identified a marginal association by capturing phenotypic similarities. These findings further demonstrate the potential of PERADIGM to identify additional genes associated with NF1-specific phenotypes beyond traditional rare variant association methods.

**Table 8:** Marginal p-values of NF1 disease for PERADIGM and SKAT-O. Each row represents a gene. Columns show the gene name, the number of variant carriers (Carrier Number), the number of carriers overlapping with NF1 diagnosis (Overlap with Disease), and the p-values calculated by PERADIGM and SKAT-O.

| Gene | Carrier Number | Overlap with Disease | P-value by PERADIGM | P-value by SKAT-O |
| --- | --- | --- | --- | --- |
| *SPRED1* | 17 | 0 | $9.15 \times 10^{-3}$ | $6.60 \times 10^{-1}$ |
| *LZTR1* | 367 | 1 | $4.76 \times 10^{-3}$ | $2.16 \times 10^{-2}$ |

## 3 Discussions

Whole exome sequencing (WES) and whole genome sequencing (WGS) have become invaluable tools for elucidating the genetic basis of human diseases, particularly rare disorders. Rare diseases, especially those with Mendelian inheritance patterns, are typically caused by pathogenic rare LoF variants in one or a few genes. Unlike common variants, rare variants often exert larger effect sizes on phenotypes, but their carriers are relatively few, even in large population cohorts such as the UK Biobank. To address the challenges of identifying these rare pathogenic variants, several rare variant association methods have been developed, including the Burden test, SKAT, and SKAT-O. These approaches improve detection power by aggregating multiple rare variants within genes or regions, making them more effective than traditional genome-wide association studies (GWAS). However, the application of these methods to rare diseases remains limited by the scarcity of affected individuals and the low frequency of causal variants. Additionally, in large biobank datasets, some individuals may have undiagnosed conditions, further reducing statistical power.

In this paper, we show that rare disease patients often exhibit distinct phenotype patterns that extend beyond their primary diagnosis. However, existing methods have not fully leveraged this comprehensive phenotypic information, focusing primarily on binary disease status. By incorporating broader phenotypic profiles, rather than restricting analyses to disease presence or absence, we may enhance our ability to detect and characterize genetic contributors to rare diseases, even in the face of limited sample sizes and missing diagnostic information.

To better utilize phenotype data, we developed PERADIGM, a framework that integrates natural language processing (NLP) techniques to extract and leverage detailed phenotype information from each individual. Unlike traditional methods that rely solely on binary disease status, PERADIGM incorporates phenotype similarity into gene identification, improving sensitivity to genetic contributors associated with disease-specific phenotypes. Through multiple case studies, we demonstrate that this approach enhances statistical power and identifies additional candidate genes that conventional methods may overlook.

We applied PERADIGM to three rare diseases: autosomal dominant polycystic kidney disease (ADPKD), Marfan syndrome, and neurofibromatosis type 1 (NF1). Each of these disorders has well-established major causative genes: *PKD1* and *PKD2* for ADPKD, *FBN1* for Marfan syndrome, and *NF1* for NF1 disease. While traditional rare variant association test methods such as the Burden test, SKAT, and SKAT-O successfully identified these primary genes in UK Biobank data, they failed to detect additional genes with smaller effects. In contrast, PERADIGM identified a broader set of genes associated with disease-specific phenotypes, some of which have supporting evidence from prior studies. To further enhance interpretability, we conducted an additional analysis to refine phenotype embeddings for diagnosed patients. Instead of embedding the full phenotype profile, we restricted embeddings to disease-related phenotypes based on expert knowledge. This targeted approach provided more explainable results by ensuring that identified genes were associated specifically with disease-relevant phenotypes rather than broader, unrelated clinical features. Our findings demonstrate that even with this stricter phenotype, PERADIGM maintained robust performance and uncovered additional associated genes. This further supports the utility of phenotype-driven gene discovery for rare diseases. For ADPKD, PERADIGM identified seven such genes, expanding beyond *PKD1* and *PKD2*. For Marfan syndrome, PERADIGM identified eight genes, significantly broadening the genetic landscape beyond *FBN1*. In the case of NF1, PERADIGM identified *NF1* as the primary gene, but marginally significant findings suggest the potential to detect additional contributors. These results illustrate PERADIGM’s ability to incorporate extensive phenotype information, leading to the identification of a more comprehensive set of genes associated with disease-specific phenotypes.

The primary innovation of PERADIGM lies in its shift from traditional binary disease-gene associations to a similarity-based framework that integrates comprehensive phenotypic information. This approach uses NLP-based embedding models to cluster individuals based on shared phenotypic patterns rather than relying on simple case-control definitions. Beyond identifying genes associated with rare LoF variant carriers, this similarity-based method can also refine disease cohort definitions and characterize individuals with distinct phenotypic profiles. In cases where patient groups are too small for traditional analyses, our method enables the identification of novel phenotype clusters, allowing for more robust investigations of disease mechanisms. By focusing on phenotypic similarity rather than individual variant effects, PERADIGM provides a powerful tool for uncovering genetic associations in rare diseases. Additionally, PERADIGM’s ability to identify genes relevant to related but distinct diseases with phenotypic overlap underscores its potential beyond rare diseases. This framework could be applied to complex traits where multiple conditions share genetic and clinical features, facilitating the discovery of novel disease subtypes and genetic modifiers.

Furthermore, PERADIGM can help infer causal relationships between genes and phenotypes by analyzing carriers’ phenotypic profiles, complementing traditional pathway-based analyses. Future improvements to the framework could involve transitioning from static embedding models like Word2Vec to context-aware models such as BERT-based architectures or large language models (LLMs). These advanced models could capture more nuanced longitudinal relationships from sequential ICD-10 code records, providing a richer representation of individual phenotypic trajectories. The success of PERADIGM highlights how integrating phenotype similarity-based embeddings into genetic research can advance our understanding of rare diseases, improve diagnostic precision, inform targeted therapies, and ultimately enhance patient outcomes.

## 4 Methods

### 4.1 UK Biobank 200K dataset and genotype data quality control

We utilized the UK Biobank 200K dataset[4] in our analysis. To mitigate potential confounding effects due to population stratification, we restricted our analysis to individuals of European ancestry. We applied several filtering criteria to ensure data quality and completeness. Specifically, we excluded individuals who lacked Hospital inpatient data or whole-exome sequencing data. After applying these filters, our final study cohort comprised 148,551 individuals of European ancestry. This refined dataset ensures a more homogeneous population with comprehensive phenotypic and genetic information, thereby enhancing the reliability and interpretability of our subsequent analyses.

To ensure the quality and reliability of the rare variant analysis, we implemented stringent quality control (QC) measures on the whole-exome sequencing (WES) data using PLINK[49]. Our QC protocol comprised several steps. We first removed variants with a minor allele frequency (MAF) exceeding 0.01, thereby retaining only rare variants for subsequent analysis. We then excluded variants that significantly deviated from Hardy-Weinberg equilibrium (HWE), with a threshold of p *<*1e-6. A significant departure from HWE can indicate genotyping errors or population stratification. Furthermore, we removed samples with missing sex information. Following these QC steps, we identified all rare predicted loss of function (pLoF) variants. These included stop-gain, stop-loss, frameshift insertion, frameshift deletion, and essential splice variants. This allowed us to focus on potentially impactful genetic alterations[50].

### 4.2 Embedding model

Embedding learning and semantic matching have a rich history in natural language processing. Mikolov et al. introduced the continuous Bag-of-Words (CBOW) and Skip-gram models, collectively known as Word2Vec, to represent words in a vector space[19]. These neural network-based models achieve state-of-the-art performance in measuring syntactic and semantic word similarity. Many studies have leveraged Word2Vec to embed medical concepts for patients using Electronic Health Record (EHR) corpora, directly utilizing textual medical records as input.

In our embedding analysis, we applied the CBOW Word2Vec model to embed the ICD-10 phenotype data from the UK Biobank. The CBOW model is an architecture used in word embedding where the surrounding context words are used to predict a target word. In CBOW, a window of context words around a target word is input to the model, and the goal is to predict the central (target) word from these context words. The model optimizes weights to maximize the probability of predicting the correct target word based on the given context. This approach is particularly well-suited for embedding ICD-10 codes based on the surrounding diagnosis records of individual patients. It effectively captures the contextual relationships between phenotypes, enabling the representation of medical concepts in a dense, continuous vector space. By leveraging these contextual relationships, the model can potentially uncover latent patterns and associations within the medical data that might not be immediately apparent through traditional analysis methods.

### 4.3 Embedding for ICD-10 codes

We aim to map phenotypes onto a static high-dimensional space, enabling numerical representation of their characteristics and facilitating similarity measurements between different phenotypes. We employed the Word2Vec embedding model to obtain static embedding vectors for the phenotypes. In the UK Biobank database, hospital inpatient data are recorded in the form of ICD-10 codes, each representing a specific disease. For instance, Q61.2 denotes, Polycystic kidney, adult type. Consequently, each patient’s record yields a sequence of ICD-10 codes, comprehensively describing their longitudinal inpatient condition. Based on this information, we developed the following approach to embedding the ICD-10 codes. For each ICD-10 code, we extracted keywords from its description, excluding punctuation and English stopwords. This process generated a vector of words for each patient based on his/her ICD-10 descriptions, with each word serving as a token in the training dataset. Each patient’s ICD-10 description was treated as a sentence. The output comprised embedding vectors for each word appearing in the inpatient dataset. We then employed an average embedding to derive the embedding vector for each ICD-10 code. This approach not only captured sequential information between different description tokens for each ICD-10 code but also leveraged detailed word tokens when two ICD-10 codes contain similar words.

### 4.4 Embedding for individual using ICD-10 codes

After obtaining the embedding vector for each ICD-10 code in the UK Biobank data, we proceeded to embed individuals using a weighted average embedding approach. This method enhances simple averaging of ICD-10 code embedding vectors by focusing only on less common ICD-10 codes (frequency ↑ 0.01) and assigning differential weights to these codes. We assigned weights by considering two pieces of information for an ICD code: (1) the significance level of each code with respect to the target disease, a process to be detailed in subsequent sections, and (2) the information content of each ICD-10 code, derived from its frequency.

For risk gene mapping, we focus on a specific disease of interest. Different phenotypes have varying degrees of associations with the target disease. Consequently, simply averaging all available ICD-10 embeddings to represent an individual’s overall phenotype embedding may be less effective to capture its relevance to the target disease. A more informative approach to characterizing an individual’s phenotypic profile in relation to a target disease involves assigning differential weights to distinct ICD-10 codes, with higher weights indicating a stronger relationship with the target disease. Target disease relevance is captured through the logistic regression model:

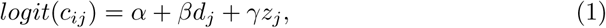

where *c_ij_* denotes the i-th ICD-10 code status of individual j, *d_j_* represents the target disease status of individual j, and *z_j_* is the vector of covariates, specifically age and sex. We conducted a comprehensive scan of all ICD-10 codes in the UK Biobank inpatient dataset for the disease of interest, calculating a p-value for each ICD-10 code. Subsequently, we utilized these p-values to derive the weight for each ICD-10 code. By employing this weighting scheme, ICD-10 codes exhibiting a more significant association with the disease of interest are assigned larger weights, thereby playing a more prominent role in the individual embedding process. We also considered each ICD-10 code’s prevalence, denoted as *r_k_*, in the embedding for each individual.

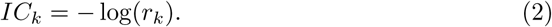

This formulation of information content assigns higher values to rarer phenotypes, reflecting our assumption that rarer phenotypes carry more information about rare diseases.

The weight *w_k_* for the *k*-th ICD-10 code is then calculated as:

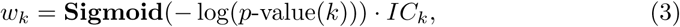

where the *p*-value represents the significance level of the *k*-th ICD-10 code in relation to the disease of interest. We use a sigmoid function to map the *p*-value onto a [0.5, 1] scale, with greater weights indicating phenotypes more relevant to the disease of interest.

The rationale behind this weighted average embedding is twofold. First, it aims to capture the differential importance of various phenotypes in relation to specific diseases, acknowledging that different phenotypes contribute distinctly to each disease. Second, it assigns more weight to rare ICD-10 codes to prevent common ICD-10 codes from overwhelming the information provided by less frequent, but potentially more informative, codes. This approach is based on the assumption that for rare diseases, the rare phenotypes each individual exhibits will carry more information about the disease. Consequently, this method ensures a more comprehensive and disease-specific representation of each individual’s phenotypic profile, potentially enhancing the detection of subtle disease associations. With such defined weights, the embedding vector for an individual *i* is calculated as:

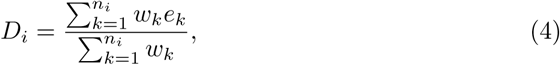

where *D_i_* is the embedding vector for individual *i*, *n_i_* is the total number of less common ICD-10 codes (frequency *↑* 0.01) recorded in individual *i*, *e_k_* is the embedding vector of the *k*-th ICD-10 code, and *w_k_* is the weight of the ICD-10 code *e_k_*. This equation shows how we represent an individual using weighted ICD-10 code embeddings.

### 4.5 Risk gene mapping

Based on the phenotype embedding of each individual, we can assign each gene a risk score for a disease of interest by investigating the similarity of the phenotype embeddings from rare LoF variant carriers with those having the target disease. A larger risk score indicates a higher likelihood that the candidate gene is associated with the disease of interest. Unlike other gene prioritization tools based on pathways or other biological criteria, we calculated the risk score solely based on phenotype similarities. This is accomplished in four steps. First, we extracted all the rare Loss of Function (LoF) variant carriers for each candidate gene, along with their genotype and phenotype information. Second, we extracted all individuals diagnosed with the target disease and their phenotype information. Third, we used the method described in the previous section to embed the LoF set carriers and disease patients, obtaining their embedding representations. Fourth, we calculated the disease risk score by comparing the phenotype similarity at the individual level between the different genes’ LoF set carriers and the diseased individuals.

Our underlying assumption is that greater similarity between disease patients and gene LoF set carriers at the phenotype embedding level indicates higher pathogenicity of the gene for the disease of interest. This is based on the hypothesis that rare LoF variants have a large effect size on the phenotype compared to common variants. Notably, we considered not only the disease status as a binary value but all relevant phenotype information related to the disease, providing a more comprehensive and reasonable approach. Finally, we ranked each gene’s risk score based on the calculated average similarity score.

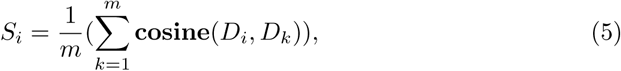

where *S_i_* is the average similarity score for individual *i* compared to the disease patients group, *m* is the number of patients in the disease of interest comparison group, *D* is the embedding vector for each individual, and the Cosine() function represents cosine similarity, commonly used in calculating the similarity between two embedding vectors. Based on the similarity score for each LoF mutation carrier, we can calculate the overall score for each candidate gene as

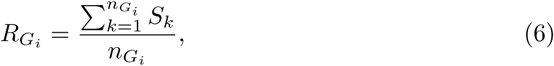

where *R_Gi_* is the risk score of gene *i* to the disease of interest, and *n_Gi_* is number of LoF carriers for gene *i*.

For each observed risk score, we employed random sampling to determine its significance level. Under the null hypothesis (*H*_0_), we derived an empirical distribution of the risk score for the target gene by repeatedly randomly selecting *n_i_*samples from the UK Biobank 200K inpatient dataset and computed the risk score for each group. This process was repeated 10,000 times to construct the empirical risk score distribution for gene *i* under the null hypothesis for the target disease.

The estimated p-value for the observed risk score is derived by calculating the Z-score using the formula:

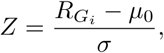

where *R_Gi_* is the observed risk score, *µ*_0_ is the average of the risk scores from randomly sampled sets of individuals, and *ϖ* is the standard deviation of the simulated risk scores. The p-value is then computed from the standard normal distribution. We applied a one-tailed test with a significance level of 0.05 and used the Bonferroni correction to control for multiple testing, assessing whether the gene’s risk score is significantly higher than expected by chance.

### 4.6 Intra-group similarity calculation

To elucidate the overall phenotype patterns within the disease group and among carriers of rare LoF variants of a candidate gene, we calculated the intra-group phenotype similarity in the embedding space. This analysis aimed to explore whether individuals within these groups share greater phenotypic similarity compared to randomly selected individuals from the UK Biobank.

First, we considered all individual pairs within the target group and calculated similarity scores for each pair within the target group. We then randomly selected an equal number of individuals from the UK WES 200K excluding the target group, and calculated pairwise similarity scores.

A difference in the similarity score box plot of the two groups would suggest that the given group exhibits a distinct intra-group phenotype pattern compared to the overall dataset. For example, this analysis could reveal whether carriers of rare LoF variants in the PKD1 gene demonstrate greater phenotypic similarity to each other than a randomly selected control group from the UK Biobank.

## Data Availability

This study did not generate any new data. All analyses were conducted using existing genotype and phenotype data from the UK Biobank, accessible at https://www. ukbiobank.ac.uk/.

## Code Availability

The code for the PERADIGM framework is publicly available at https://github.com/JJJJJasonZheng/PERADIGM.

