## Supplement for "PERADIGM: Phenotype Embedding Similarity-based Rare Disease Gene Mapping"

#### Supplementary Materials

**Wangjie Zheng**[1], **Yuhan Xie**[1], **Jianlei Gu**[1], **Hongyu Li**[1], **Stefan Somlo**[2], **Whitney Besse**[2], **Hongyu Zhao**[1,*]

[1]Department of Biostatistics, Yale University, New Haven, CT 06510, United States
[2]Department of Internal Medicine, Section of Nephrology, Yale School of Medicine, New Haven, CT 06510, United States

### Appendix A Supplementary Materials

#### A.1 Genotype quality control results

Table A1 below shows the number of variants on each chromosome in different stages: raw genotype data, after quality control, and LoF variants.

| Chromosome | Raw Variants | After QC | LoF Variants |
|---|---|---|---|
| Chr1 | 1,783,906 | 1,712,567 | 45,456 |
| Chr2 | 1,310,712 | 1,257,244 | 31,833 |
| Chr3 | 1,039,941 | 998,606 | 26,444 |
| Chr4 | 717,783 | 685,611 | 18,004 |
| Chr5 | 790,760 | 757,766 | 19,752 |
| Chr6 | 887,003 | 847,539 | 23,231 |
| Chr7 | 858,981 | 821,664 | 21,040 |
| Chr8 | 654,090 | 627,300 | 15,079 |
| Chr9 | 774,556 | 742,399 | 17,595 |
| Chr10 | 733,427 | 703,066 | 17,526 |
| Chr11 | 1,059,628 | 1,019,090 | 26,958 |
| Chr12 | 950,532 | 910,028 | 22,992 |
| Chr13 | 320,631 | 306,289 | 8,361 |
| Chr14 | 557,237 | 533,269 | 13,266 |
| Chr15 | 620,095 | 594,231 | 15,532 |
| Chr16 | 875,192 | 841,107 | 19,373 |
| Chr17 | 1,047,224 | 1,006,046 | 25,414 |
| Chr18 | 286,694 | 274,147 | 6,798 |
| Chr19 | 1,210,652 | 1,156,885 | 32,429 |
| Chr20 | 459,900 | 441,748 | 10,394 |
| Chr21 | 195,726 | 186,232 | 4,520 |
| Chr22 | 414,980 | 397,963 | 9,207 |
| **Total** | 17,549,650 | 16,820,797 | 431,204 |

**Table A1**: **Summary of Variants After QC and LoF Annotation.** Each row represents a chromosome. Columns show the number of variants before quality control (Raw Variants), after quality control (After QC), and the number of annotated loss-of-function (LoF) variants. The final row reports the totals across all chromosomes.

#### A.2 P-Value inflation analysis of PERADIGM

To evaluate the behavior of test statistics under the null hypothesis, we investigated possible p-value inflation in PERADIGM by calculating the genomic inflation factor ($\lambda$) for three representative conditions: Marfan syndrome, ADPKD, and NF1 disease. The genomic inflation factor is defined as the ratio of the median of the observed test statistics to the expected median under the null hypothesis. A $\lambda$ value greater than 1 indicates potential p-value inflation (which can result in false positives), while $\lambda$ values less than 1 suggest p-value deflation (overly conservative results). A value of $\lambda = 1$ reflects an ideal behavior with no inflation or deflation of test statistics.

The results of our analysis show that the $\lambda$ values for ADPKD (0.9904), Marfan syndrome (0.9709) and NF1 (0.9838) closely align with the expected value of 1. This indicates that PERADIGM does not exhibit p-value inflation and demonstrates that the test statistics under the null hypothesis behave as expected. These results provide confidence in the reliability of the p-values produced by PERADIGM for subsequent statistical analyses.

#### A.3 Comparison of disease and non-disease group risk score distribution

Our analysis compared the risk scores between disease and non-disease groups for three rare disorders: ADPKD, Marfan syndrome, and NF1 disease. For all three conditions, the disease groups exhibited significantly higher risk scores compared to their respective non-disease counterparts (random sample with equal size as the disease group). This marked difference in risk scores demonstrates the method's robust ability to differentiate between affected and unaffected individuals. These findings underscore the potential of our risk score approach as a valuable tool for distinguishing disease status in rare Mendelian disorders, even within large-scale genomic datasets where such conditions are infrequently represented.

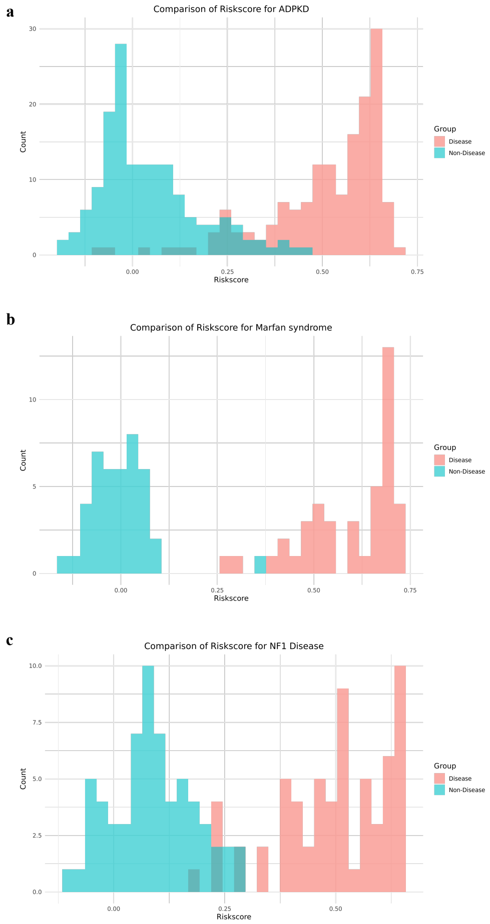


**Fig. A1**: **Comparison of disease and non-disease group risk scores. a.** Histogram of gene risk scores for individuals diagnosed with ADPKD and controls. **b.** Histogram of gene risk scores for individuals diagnosed with Marfan syndrome and controls. **c.** Histogram of gene risk scores for individuals diagnosed with NF1 and controls.

#### A.4 Histogram of intra-group similarity score between diseases and gene rare LoF carriers

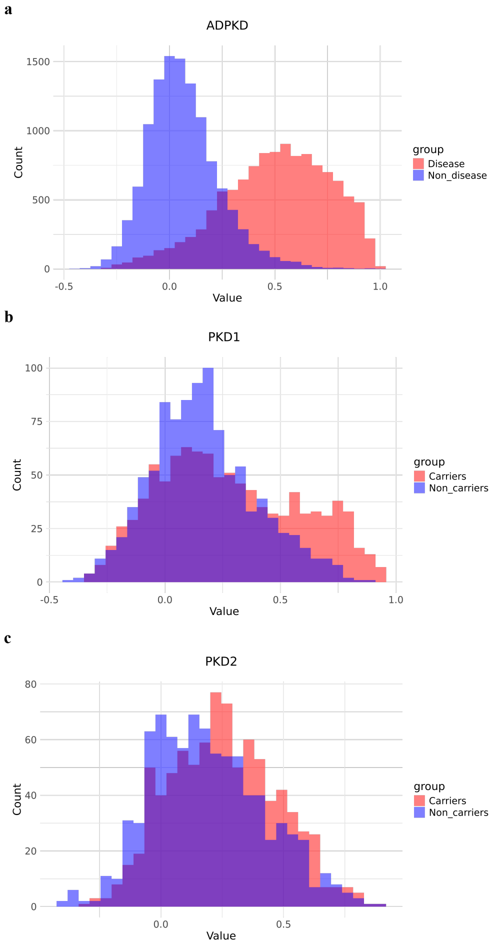


**Fig. A2**: **Histogram of pairwise similarity score analysis for ADPKD. a.** Distribution of similarity scores among ADPKD patients compared to non-disease individuals. **b.** Distribution of similarity scores for individuals carrying rare *PKD1* LoF variants compared to non-carriers. **c.** Distribution of similarity scores for individuals carrying rare *PKD2* LoF variants compared to non-carriers.

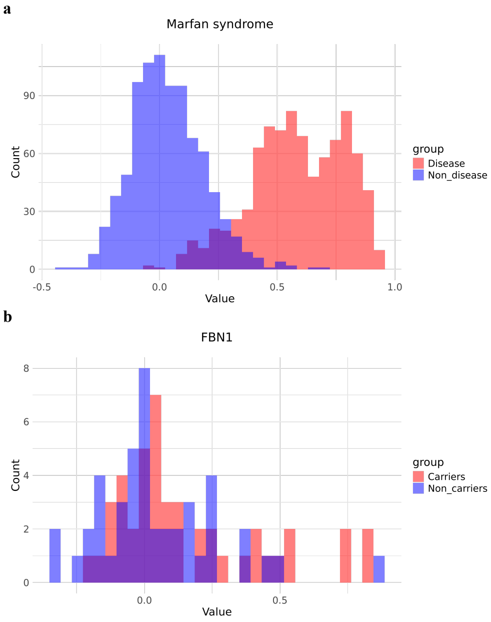


**Fig. A3**: **Histogram of pairwise similarity score analysis for Marfan syndrome. a.** Distribution of similarity scores among Marfan syndrome patients compared to non-disease individuals. **b.** Distribution of similarity scores for individuals carrying rare *FBN1* LoF variants compared to non-carriers.

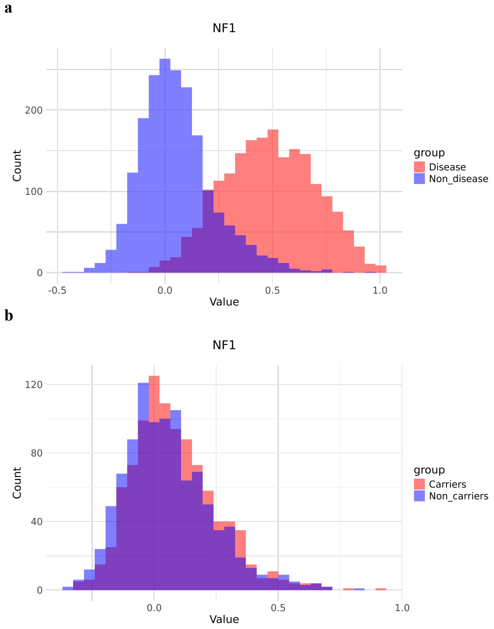


**Fig. A4**: **Histogram of pairwise similarity score analysis for NF1. a.** Distribution of similarity scores among NF1 patients compared to non-disease individuals. **b.** Distribution of similarity scores for individuals carrying rare *NF1* LoF variants compared to non-carriers.

#### A.5 Violin plot of risk scores for the significant genes

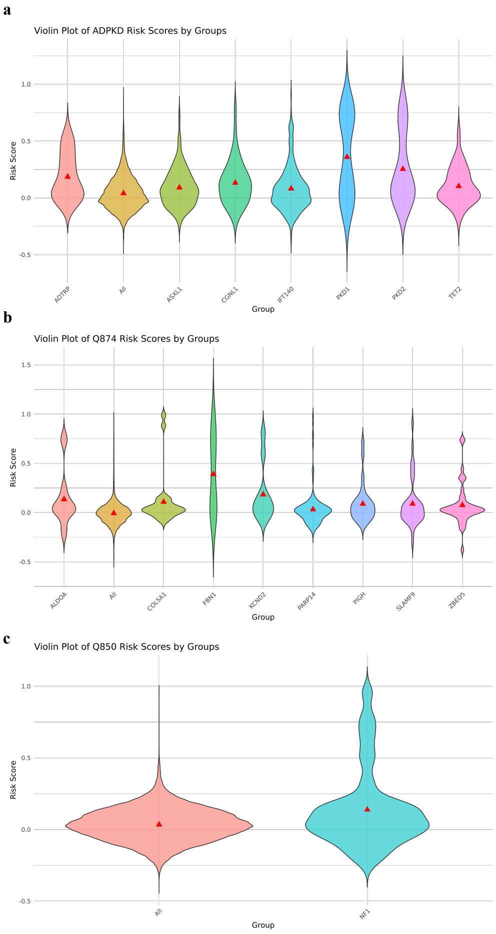


**Fig. A5**: **Violin plot of risk scores by groups. a.** Distribution of risk scores for individuals carrying rare LoF variants in genes significantly associated with ADPKD. The "All" group represents the risk score distribution of all individuals in the dataset with respect to ADPKD-specific phenotypes. **b.** Distribution of risk scores for individuals carrying rare LoF variants in genes significantly associated with Marfan syndrome. The "All" group reflects the risk score distribution for the entire cohort based on Marfan syndrome-specific phenotypes. **c.** Distribution of risk scores for individuals carrying rare LoF variants in genes significantly associated with NF1 disease. The "All" group shows the risk score distribution across all individuals with respect to NF1-specific phenotypes.
